# Effect of analytical variability in estimating EEG-based functional connectivity

**DOI:** 10.1101/2023.08.17.553675

**Authors:** Sahar Allouch, Aya Kabbara, Joan Duprez, Véronique Paban, Mohamad Khalil, Julien Modolo, Mahmoud Hassan

## Abstract

The significant degree of variability and flexibility in neuroimaging analysis approaches has recently raised concerns. When running any neuroimaging study, the researcher is faced with a large number of methodological choices, often made arbitrarily. This can produce substantial variability in the results, ultimately hindering research replicability, and thus, robust conclusions. Here, we addressed the analytical variability in the EEG source connectivity pipeline and its effects on outcomes consistency. Like most neuroimaging analyses, the EEG source connectivity analysis involves the processing of high-dimensional data and is characterized by a complex workflow that leads to high analytical variability. In this study, we focused on source functional connectivity variability induced by three key factors along the analysis pipeline: 1) number of EEG electrodes, 2) inverse solution algorithms, and 3) functional connectivity metrics. Outcomes variability was assessed in terms of group-level consistency, inter-, and intra-subjects similarity, using resting-state EEG data (n = 88). As expected, our results showed that different choices related to the number of electrodes, source reconstruction algorithm, and functional connectivity measure substantially affect group-level consistency, between-, and within-subjects similarity. We believe that the significant impact of such methodological variability represents a critical issue for neuroimaging studies that should be prioritized.

**Highlights:**

- The significant impact of methodological variability is a recognized critical priority issue for neuroimaging studies.
- Analytical variability related to the number of electrodes, source reconstruction algorithm, and functional connectivity measure is a prominent issue in the EEG source connectivity analysis.
- Group-level consistency, between-, and within-subjects similarity are substantially affected by analytical variability in the EEG source connectivity analysis.

## 1. Introduction

### 1.1. Analytical variability in neuroimaging research

In the course of running a study, the researcher is faced with a large number of choices often made arbitrarily. Those choices are known as the researcher’s “degree of freedom” (Simmons, Nelson, and Simonsohn 2011), and are spread out across all research phases (Wicherts et al. 2016). Recently, the question of analytical flexibility has received much attention in many scientific fields as a potential source of poor reproducibility (Botvinik-Nezer et al. 2020; Wicherts et al. 2016; Simmons, Nelson, and Simonsohn 2011; Silberzahn et al. 2018; Carp 2012). The risks of high variability in analytical approaches are twofold. First, one may chase statistical significance by taking advantage of the researcher’s degrees of freedom, i.e., run several analyses and modify the study while progressing until finding a significant positive result, and then only reporting the positive result and the corresponding analysis (Bishop et al. 2015; Simmons, Nelson, and Simonsohn 2011; Carp 2012). Second, interestingly, the researcher may run only one analysis, and that analysis yields significant positive results, but only due to certain choices along the analysis pipeline that are not always justified or documented. Hence, analytic decisions made without direct attempts to achieve statistical significance can still produce high variability in results (Simmons, Nelson, and Simonsohn 2011; Carp 2012; Silberzahn et al. 2018; Botvinik-Nezer et al. 2020), thereby affecting conclusions that could be drawn from a study (Botvinik-Nezer et al., 2020), and ultimately hindering the reproducibility of scientific research (Wicherts et al., 2016).

Due to the high dimensionality of the data and increasing complexity of analysis workflows, analytical variability issues are thought to be prominent in neuroimaging research (Botvinik-Nezer et al. 2020). Recently, those issues were addressed in the field of fMRI (Botvinik-Nezer et al. 2020) and EEG (https://www.eegmanypipelines.org) by large collaborative initiatives where independent teams were asked to analyze the same data set and test the same hypotheses. Moreover, the sensitivity of EEG results to the variability in preprocessing strategies was investigated in (Šoškić et al. 2022; Clayson et al. 2021; Robbins et al. 2020). Study outcomes were mainly affected by the high-pass filter cut-off, artifact removal method, baseline duration, reference, measurement latency and locations, and amplitude measure (peak vs. mean). Different analysis software (Bowring, Maumet, and Nichols 2019; Glatard et al. 2015), software versions (Gronenschild et al. 2012) and operating systems (Glatard et al. 2015; Gronenschild et al. 2012) were also studied as substantial sources of variability in fMRI preprocessing. Similarly, (Aya Kabbara et al. 2022) revealed that a considerable variability in results is observed when using different software tools to preprocess EEG signals.

### 1.2. Analytical variability in EEG source connectivity analysis

In this work, we are particularly interested in the question of analytical variability in the specific context of functional brain connectivity inferred from EEG data (Hassan and Wendling 2018; Hassan et al. 2015, 2014; A. Kabbara et al. 2017; Mehrkanoon et al. 2014). Mainly, the EEG source connectivity analysis consists of two main steps: (i) reconstructing the dynamics of cortical sources by solving the so-called EEG inverse problem, and (ii) computing functional connectivity between the reconstructed sources. Although these two steps are relatively standard in the field of EEG-based brain connectivity, combining them can be much more cumbersome than it seems in terms of choices and flexibility. Dozens of substeps are to be made, each entailing numerous choices that are often arbitrary. For instance, during the first steps of the experiment, the spatial density (i.e., number of electrodes) of the EEG system must be determined. When solving the inverse problem, a large set of mathematical methods is available, each imposing assumptions and constraints regarding the spatial and temporal properties of the reconstructed sources, without mentioning the various parameters that require tuning for each of those algorithms. This is also valid for connectivity measures, with metrics assessing various signal features such as phase synchronization, amplitude synchronization, omitting or keeping zero-lag connections, and a set of parameters that require tuning in each method. Thus, the numerous choices to be made in the EEG source connectivity analysis pipeline have the potential to be problematic, which has not been fully explored to date. The researcher’s high degree of freedom could produce a substantial variability in the reported results, ultimately hindering replicability in EEG source connectivity research. Therefore, the overall goal of the present paper is to call into question the analytical variability in the EEG source connectivity pipeline and its effects on the consistency/discrepancy of the outcomes. Mainly, we focused on three key factors along the analysis:

#### i. The number of EEG electrodes (i.e., electrode density)

The spatial density of standard EEG systems ranges from 19 to 256 sensors and several studies investigated its effect on EEG source localization in simulations in the context of epilepsy (Song et al. 2015; Sohrabpour et al. 2015; Lantz et al. 2003). More specifically, it has been shown that a higher number of electrodes is associated with a significant decrease in localization error (Lantz et al. 2003; Song et al. 2015; Sohrabpour et al. 2015). In two previous simulation studies (Allouch et al. 2022, 2023), we showed a significant effect of the number of electrodes (19, 32, 64, 128, 256) on the accuracy of reconstructed cortical networks and their correspondence with the reference simulated networks. In this article, we wanted to investigate if this effect stands in the experimental EEG context. Thus we tested three electrode configurations (64, 32, 19 channels; Higher densities were not investigated due to the dataset characteristics: 64-channels EEG system was used) on the outcomes’ variability.

#### ii. The source reconstruction algorithms

Another critical influencing factor in the EEG source connectivity analysis is the algorithm chosen to reconstruct cortical source activity. Several studies quantifying the performance of different inverse methods, in simulated and experimental EEG/MEG data, concluded that the choice of the inverse method significantly influences source estimation results (Anzolin et al. 2019; Mahjoory et al. 2017; Hedrich et al. 2017; Bradley et al. 2016; Grova et al. 2006; Halder et al. 2019; Tait et al. 2021; Allouch et al. 2022, 2023). In this article, we choose to test three widely used source reconstruction algorithms, specifically, eLORETA, LCMV, and wMNE.

#### iii. The functional connectivity metrics

The choice of the functional connectivity metric is also a critical step when reconstructing brain networks. A wide range of measures are used in the field, and each differs in the aspect of the data under investigation (linear/non-linear, amplitude/phase-synchronization, time/spectral domain, prone/robust to source leakage, etc…), (see (Friston 2011; Pereda, Quiroga, and Bhattacharya 2005; Cao et al. 2022) for a review), resulting in significant variability of performance and interpretations (Colclough et al. 2016; H. E. Wang et al. 2014; Wendling et al. 2009; Hassan et al. 2017; Allouch et al. 2022, 2023). Here, the tested measures included: PLV, AEC, PLI, wPLI, and corrected AEC and PLV, since they are widely used in functional connectivity studies, and cover qualitatively different approaches (e.g., phase-based *versus* amplitude-based).

## 2. Materials and Methods

The full pipeline of our study is summarized in Fig. 1.

### 2.1. Dataset

#### 2.1.1. Data acquisition

125 healthy subjects (78 Females) were recruited at Aix Marseille University, France. Their age ranged between 20 and 75 years old. (mean = 42, sd = 16). The study was approved by the “Comité de Protection des Personnes” (agreement n° 19.09.12.44636). Some of the data was used in previous studies (A. Kabbara, Paban, and Hassan 2021; Aya Kabbara et al. 2020; Paban et al. 2019). Participants were asked to relax for 5 minutes with their eyes closed without falling asleep. EEG data were collected using a 64-channel Biosemi ActiveTwo system (Biosemi Instruments, Amsterdam, The Netherlands) positioned according to the standard 10–20 system montage. Two bilateral electrooculogram electrodes (EOG) recorded horizontal movements. Electrode impedances were kept below 20 *k*Ω. EEG data were originally sampled at 1024 or 2048 Hz.

**Fig. 1.**
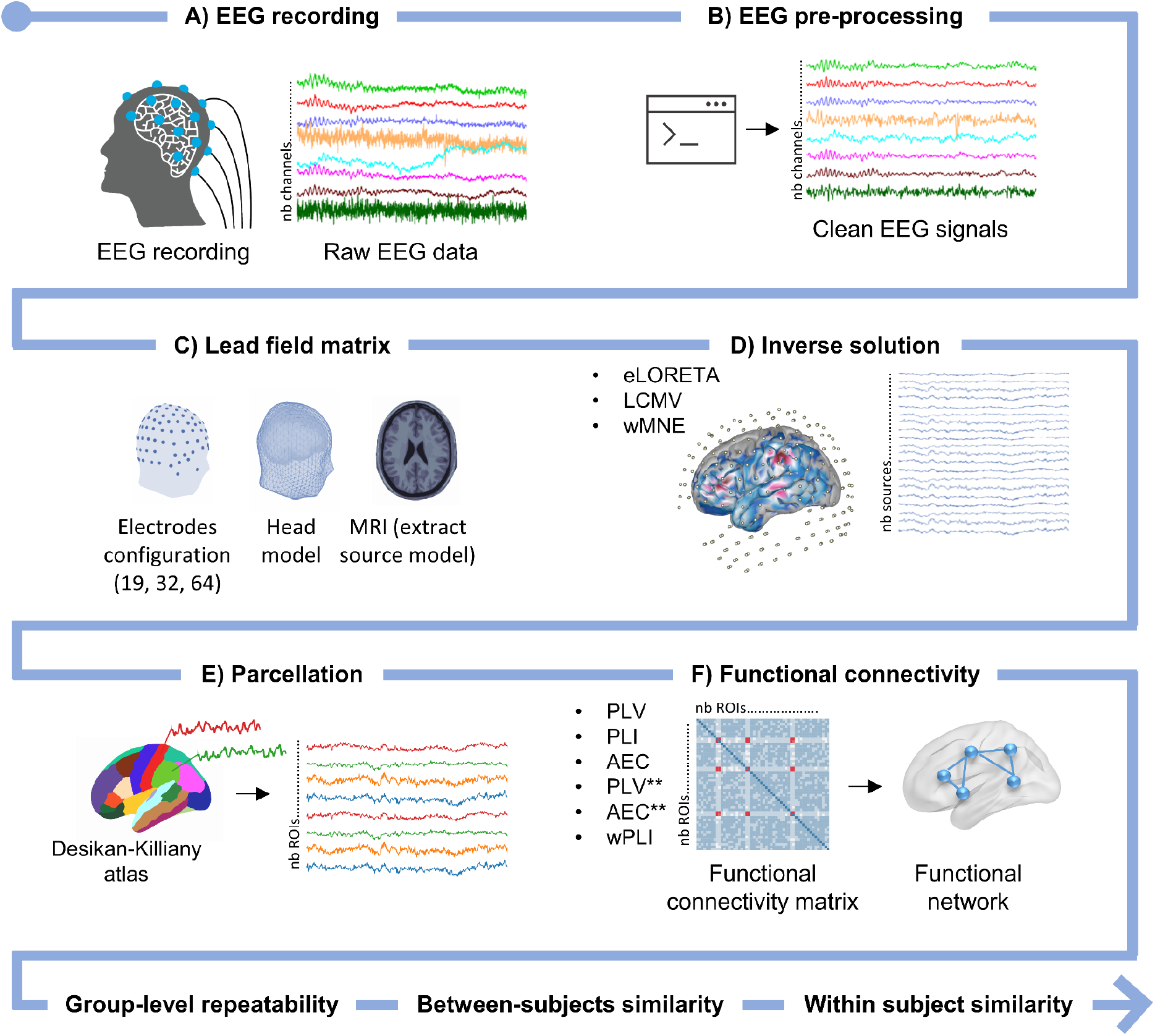
Full pipeline of the study. The EEG source connectivity analysis consisted of A) scalp EEG recordings; B) EEG signals preprocessing; C) Gain (lead field) matrix computation; D) solving the inverse problem to reconstruct cortical activity; E) regional time series extraction; F) functional connectivity computation. Networks reconstructed using different electrode configurations, inverse solution algorithms, and connectivity measures were compared based on group-level repeatability, between-subjects similarity, and within-subject similarity. *eLORETA - exact low-resolution electromagnetic tomography. LCMV - linearly constrained minimum norm beamforming. wMNE - weighted minimum norm estimate. PLV - phase-locking value. PLI - phase-lag index. AEC - amplitude envelope correlation. wPLI - weighted phase-lag index. Asterisks (**) - source leakage correction*.

#### 2.1.2. Data preprocessing

Data were re-sampled to a common frequency of 512 Hz and segmented into 10-seconds epochs. Preprocessing was done using Automagic Matlab toolbox (Pedroni, Bahreini, and Langer 2019) (https://github.com/methlabUZH/automagic). Automagic preprocessing parameters are detailed in the Supplementary Materials. The number of interpolated channels did not exceed 15% of the total number of channels; otherwise, the epoch was rejected. For each subject, we chose the best 30 epochs based on Automagic quality metrics (please refer to Supplementary Materials). Subjects having less than 30 clean epochs were excluded from analysis. After preprocessing, 88 subjects were included in the analysis. Finally, the 64-channel montage wass subsampled into 32- and 19-channel montages.

### 2.2. EEG source connectivity

As aforementioned, the “EEG source connectivity” method refers to the assessment of functional connectivity between cortical sources. It includes two main steps: 1) reconstructing the temporal dynamics of cortical sources by solving the EEG inverse problem, and 2) assessing the functional connectivity between the reconstructed sources (Hassan and Wendling 2018; Hassan et al. 2015, 2014; A. Kabbara et al. 2017; Mehrkanoon et al. 2014).

#### 2.2.1 EEG inverse solution

The source model, which provides information about the location and orientation of the dipole sources to be estimated, was computed based on ICBM152 MRI template using the Boundary Element Method (BEM) in OpenMEEG (Gramfort et al. 2010) implemented in Fieldtrip toolbox (Oostenveld et al. 2011). EEG signals *X*(*t*) recorded from *Q* channels (64, 32, or 19 channels) can therefore be expressed as a linear combination of *P* time-varying current dipole sources *S*(*t*):

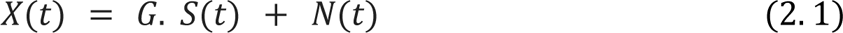

where *G* (*Q* × *P*) is the lead field (gain) matrix and *N*(*t*) is the additive noise. *G* reflects the contribution of each cortical source to scalp sensors, and is computed from a multiple-layer head model and the position of the *Q* electrodes. In order to assess the effect of sensors’spatial resolution on the reconstructed cortical networks, the lead field matrix was computed using different electrode montages (64, 32, and 19 channels). The source model was constrained to a field of current dipoles homogeneously dispersed over the cortex and normal to the cortical surface. In this case, the inverse problem can be reduced to computing the magnitude of dipolar sources 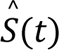 as follows:

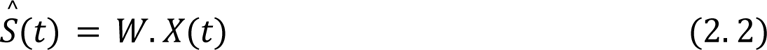

Since the problem is ill-posed (undetermined, the number of sources being far greater than the number of observations), mathematical and/or biophysical or electrophysiological assumptions need to be imposed to compute *W* and find a unique solution that fits the data (see (Awan, Saleem, and Kiran 2019; Grech et al. 2008; Baillet, Mosher, and Leahy 2001) for a review).

In this study, we chose to test the variability in the networks obtained using three different source reconstruction algorithms:

##### i. Weighted minimum norm estimate (wMNE)

The minimum norm estimate, initially proposed by (Hämäläinen and Ilmoniemi 1994), and widely used in EEG source imaging, searches for a solution that fits the data while minimizing the energy (minimum least-square error, L2-norm). An intrinsic consequence of this constraint is a bias toward superficial sources generating strong fields with less energy due to their proximity to electrodes (He et al. 2018; Michel and Brunet 2019). To compensate for the tendency of MNE to favor weak and surface sources, the weighted minimum norm estimate (wMNE) (Lin et al. 2006; Fuchs et al. 1999) sets the diagonal elements of *B* (eq. 2.4) inversely proportional to the norm of the lead field vectors, essentially assuming a priori that sources that only weakly influence M/EEG must have a higher variance to be measured by sensors (Tait et al. 2021).

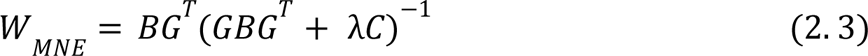

where λ is the regularization parameter, and *C* the noise covariance matrix.

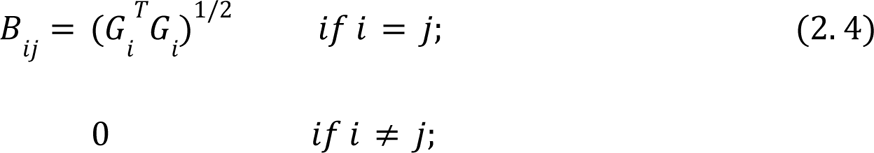

##### ii. Exact low-resolution brain electromagnetic tomography (eLORETA)

eLORETA belongs to the family of weighted minimum norm inverse solutions. In addition to compensating for depth bias, it also has exact-zero error localization in the presence of measurement and structured biological noise (Pascual-Marqui 2007):

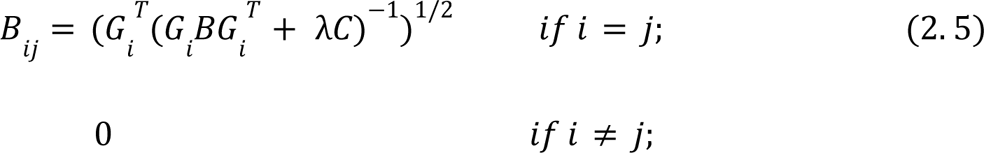

##### iii. Linearly constrained minimum-variance (LCMV) beamformer

Beamformers (a.k.a. spatial filters or virtual sensors), originally established for radar and sonar signal processing, are now widely used in source imaging, mainly with MEG data (Michel et al. 2004). The basic idea of beamformer approaches is to discriminate between signals arriving from a location of interest and those originating elsewhere (Baillet, Mosher, and Leahy 2001). Specifically, the LCMV beamformer (Van Veen et al. 1997) estimates the activity for a source at a given location while simultaneously suppressing (i.e., setting null values) contributions from all other sources and noise captured in the data covariance.

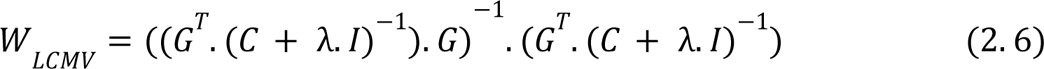

The estimation of matrix *W* were done on a high-resolution surface mesh (15000 vertices). Spatially close brain sources were then clustered (average signal) into 68 regional time series *R*(*t*) based on the regions of interest (ROIs) of the Desikan-Killiany atlas (Desikan et al. 2006).

#### 2.2.2. Functional connectivity

The next step following the reconstruction of cortical dynamics, is to assess functional connectivity, i.e. statistical interdependence between spatially distant brain regions (Friston 2011). As aforementioned, a plethora of methods are proposed in the literature which are either linear or nonlinear, parametric or nonparametric, based on phase and/or amplitude synchronization, computed in time and/or frequency domain, robust or prone to source leakage (see (Friston 2011; Pereda, Quiroga, and Bhattacharya 2005; Cao et al. 2022) for a review). At the end of this step, an *R* × *R* matrix is obtained, where each entry *a_ij_* of the matrix represents the weight of the connection linking node (i.e. ROI) *i* to node *j*. Since our intent was not to present an exhaustive comparison between all available metrics, our study covered a total of six frequently used metrics while representing the main types of approaches:

##### i. Phase-locking value (PLV)

For two signals *x*(*t*) and *y*(*t*), the phase-locking value (Lachaux et al. 2000) is defined as:

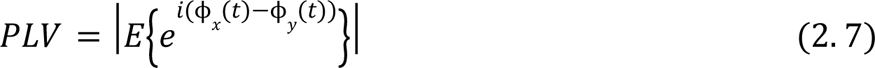

where *E*{.} is the expected value operator and ϕ(*t*) is the instantaneous phase derived from the Hilbert transform.

##### ii. Phase-lag index (PLI)

The PLI, originally proposed in (Stam, Nolte, and Daffertshofer 2007), is a measure of the asymmetry of the distribution of phase differences between two signals. It aims at overcoming the issue of source leakage by discarding phase differences centered around 0 and π, i.e., removing zero-lag connections. It is an estimation of the extent of non-equiprobability of phase leads and lags between signals (Vinck et al. 2011). For two signals *x*(*t*) and *y*(*t*), PLI is defined as follows:

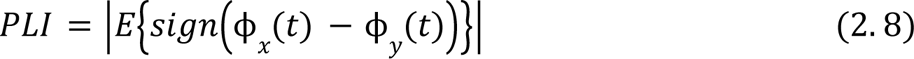

where *E*{.} is the expected value operator, and ϕ(*t*) is the instantaneous phase derived from the Hilbert transform.

##### iii. Weighted phase-lag index (wPLI)

The weighted phase-lag index attempts to further weight the metric away from zero-lag contributions (Vinck et al. 2011). The contribution of observed phase leads and lags is weighted by the magnitude of the imaginary component of the cross-spectrum. This results in reduced sensitivity to additional, uncorrelated noise sources and increased statistical power to detect changes in phase-synchronization. wPLI is defined as follows:

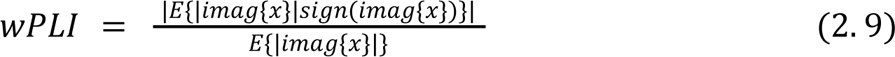

Where *imag*{*x*} denotes the imaginary part of the signal’s cross-spectrum.

##### iv. Amplitude envelope correlation (AEC)

AEC denotes the Pearson correlation between the signals’ envelopes derived from the Hilbert transform (Hipp et al. 2012; Brookes et al. 2011).

##### v. PLV and AEC with source leakage correction (PLV**, AEC**)

We computed PLV and AEC between time courses corrected for source leakage. Zero-lag signal overlaps are removed by regressing out (orthogonalizing with respect to) the linear projection of the regional time course (Brookes, Woolrich, and Barnes 2012). Since orthogonalization can be done in two directions (*x* to *y*, *y* to *x*), we computed PLV and AEC for both directions of orthogonalized time series and then averaged the obtained values. We also tested a multivariate symmetric orthogonalization approach (Results are shown in the Supplementary Materials): the closest orthonormal matrix to the uncorrected regional time courses was first computed; then the magnitudes of the orthogonalized vectors were adjusted iteratively to minimize least-squares distances between corrected and uncorrected signals (Colclough et al. 2015).

### 2.3. Results quantification

The effects of the different analytical choices (number of electrodes, source reconstruction algorithm, functional connectivity measure) were estimated by assessing the:

i. Group-level consistency/repeatability,
ii. Between-subjects similarity/variability,
iii. Within-subject similarity/variability.

#### 2.3.1. Group-level consistency

A split-half reliability approach was used to assess group-level consistency (Colclough et al. 2016). The dataset (88 subjects) was randomly divided into two groups *G*1 and *G*2. Group connectivity matrices (**C_G1_** and **C_G2_**) were obtained by averaging connectivity matrices across all subjects within a single group. Then, Pearson correlation was computed between averaged matrices **C_G1_** and **C_G2_**. This was repeated for 100 iterations.

To investigate the effect of source leakage on group-level consistency, we computed the correlation between edge length (i.e., distance between ROIs) and its contribution to group-level consistency. The contribution of individual edges (Finn et al. 2015; Colclough et al. 2016) to the group-level consistency was defined as the element-wise product between edge vectors of matrices **C_G1_** and **C_G2_** (after z-score normalization). The contribution of individual edges was averaged across all iterations and correlated with the distance separating distinct ROIs.

#### 2.3.2. Between-subjects similarity

We computed the Pearson correlation between the connectivity matrices of the different subjects in the dataset (Colclough et al. 2016). The distribution of correlation values reflects the between-subjects similarity/variability obtained for different channel densities, inverse solutions, and connectivity measures.

#### 2.3.3. Within-subject similarity

Pearson correlation was computed between all connectivity matrices of the different epochs within a single subject (30 epochs). We then plotted the distribution of averaged correlation values within each subject. Higher correlation values reflect thus an intra-subject similarity, that is, consistency across all epochs of the subject.

#### 2.3.4. Statistical Testing

All statistical analyses were performed using R (R Core Team 2020). Generalized linear models were used to assess the effect of (i) number of electrodes, (ii) inverse solution, and (iii) functional connectivity metric as fixed effects on group-level repeatability and between-subjects similarity. Beta regression model was used to assess the effects of different tested factors on the absolute value of the within-subjects similarity. These models were chosen after careful inspection of the model’s assumptions based on visual inspections of the model’s residuals distributions. In each case, the model that best met these assumptions was chosen. *Post-hoc* analyses were performed using z-tests with the *glht* function of the *multcomp* package (Hothorn, Bretz, and Westfall 2008) that provides corrected *p*-values. Models’ R^2^ were calculated with the *r.squaredGLMM* function of the *MuMin* package (Barton 2009) to estimate the variance explained by models. The significance threshold was set at its usual value of 0.05.

## 3. Results

Violin plots of the distributions of group level consistency, between-, and within-subject similarity are shown in Fig. 2, Fig. 3, and Fig. 4, respectively. The results reported here correspond to the alpha ([8-13 Hz]) frequency band, whereas results obtained for other frequency bands are shown in Fig. S1-S3 in the Supplementary Materials.

**Fig. 2.**
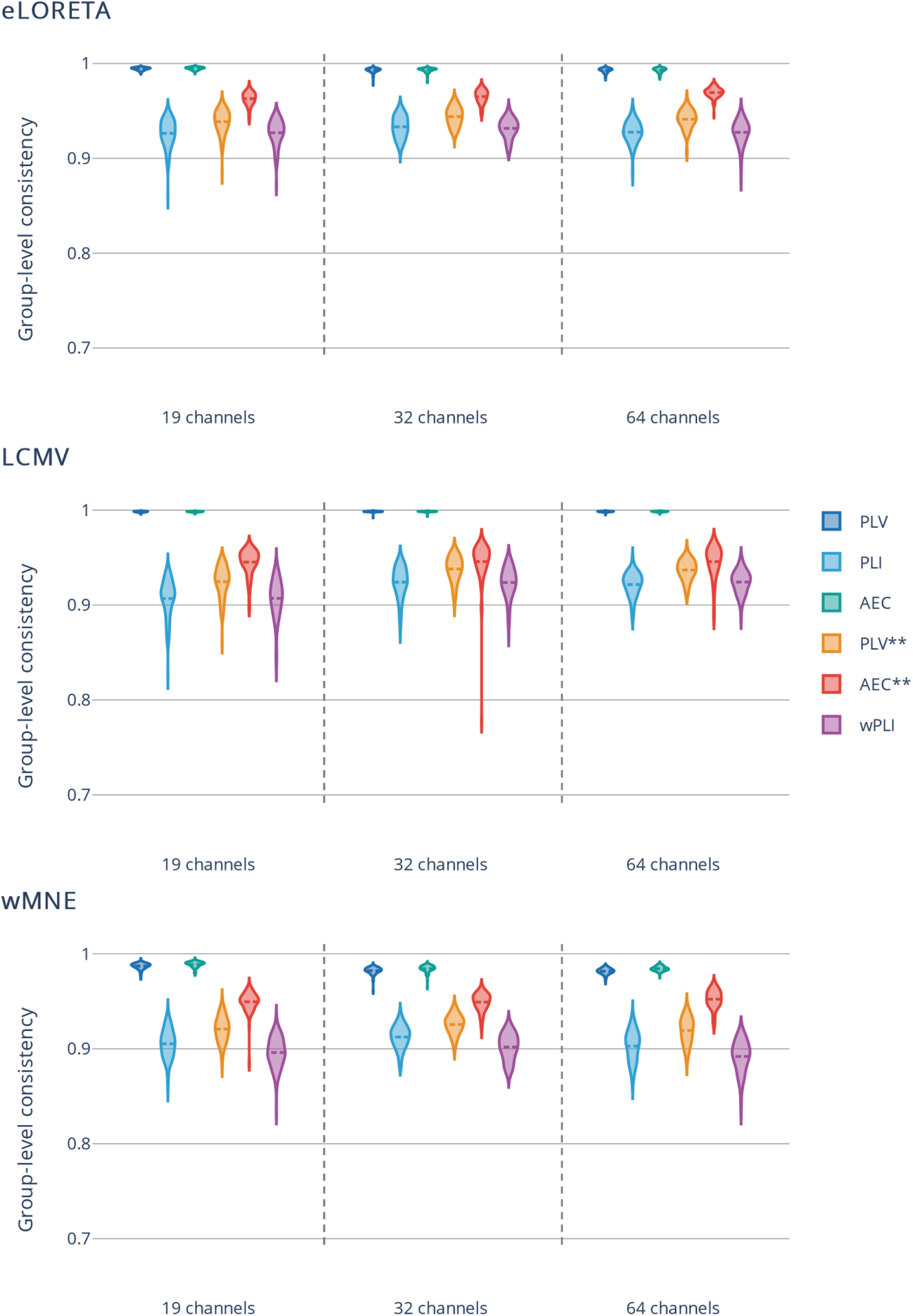
Group-level consistency. The dataset was divided into two halves 100 times. For each condition (number of electrodes x inverse solution algorithm x connectivity measure), the Pearson correlation was computed between averaged connectivity matrices inferred from separate halves of the dataset. *eLORETA - exact low-resolution electromagnetic tomography. LCMV - linearly constrained minimum norm beamforming. wMNE - weighted minimum norm estimate. PLV - phase-locking value. PLI - phase-lag index. AEC - amplitude envelope correlation. wPLI - weighted phase-lag index. Asterisks (**) - source leakage correction*.

**Fig. 3.**
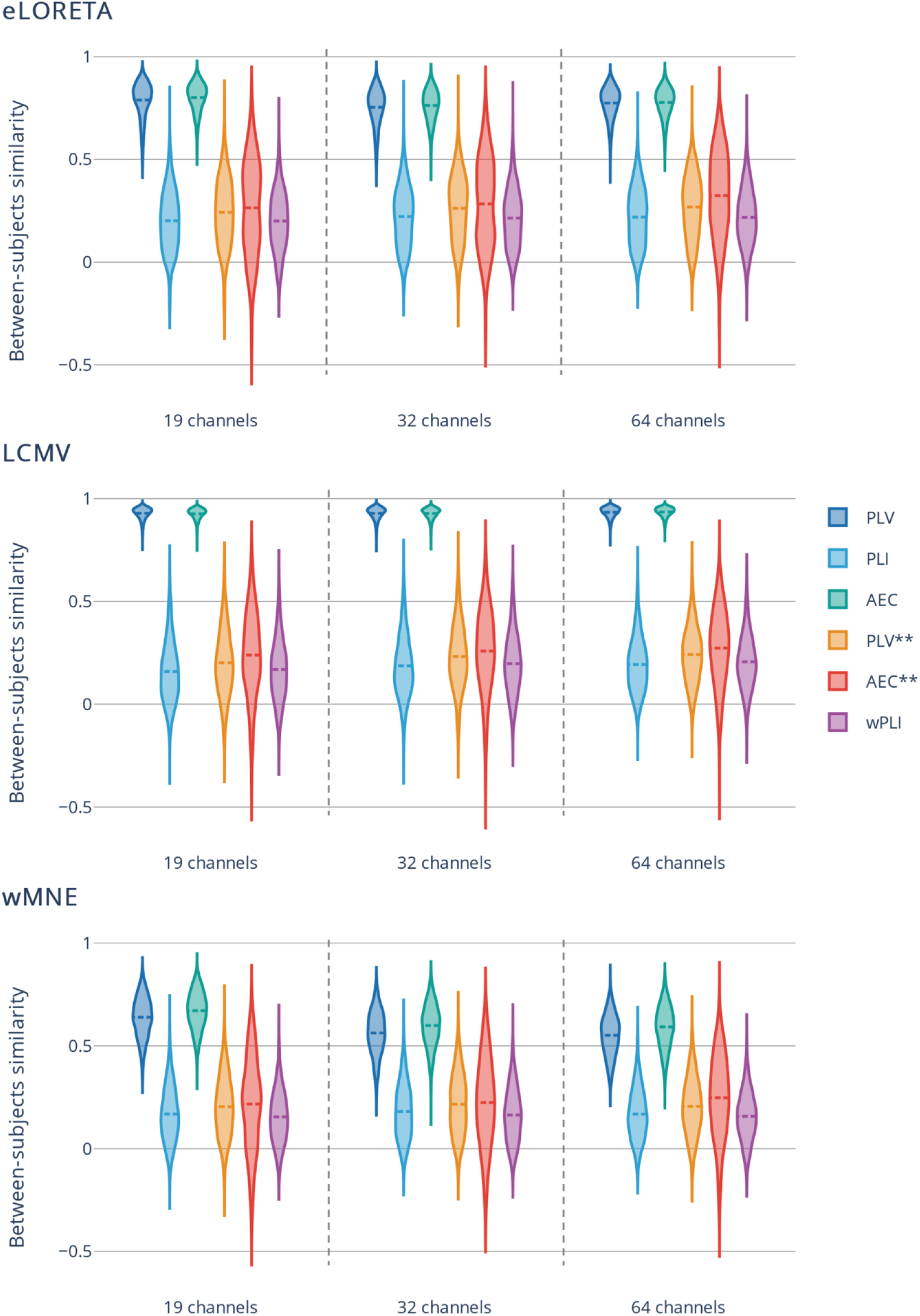
Between-subjects similarity. Pearson correlation was computed between all subjects for each condition (number of electrodes x inverse solution algorithm x connectivity measure). *eLORETA - exact low-resolution electromagnetic tomography. LCMV - linearly constrained minimum norm beamforming. wMNE - weighted minimum norm estimate. PLV - phase-locking value. PLI - phase-lag index. AEC - amplitude envelope correlation. wPLI - weighted phase-lag index. Asterisks (**) - source leakage correction*.

**Fig. 4.**
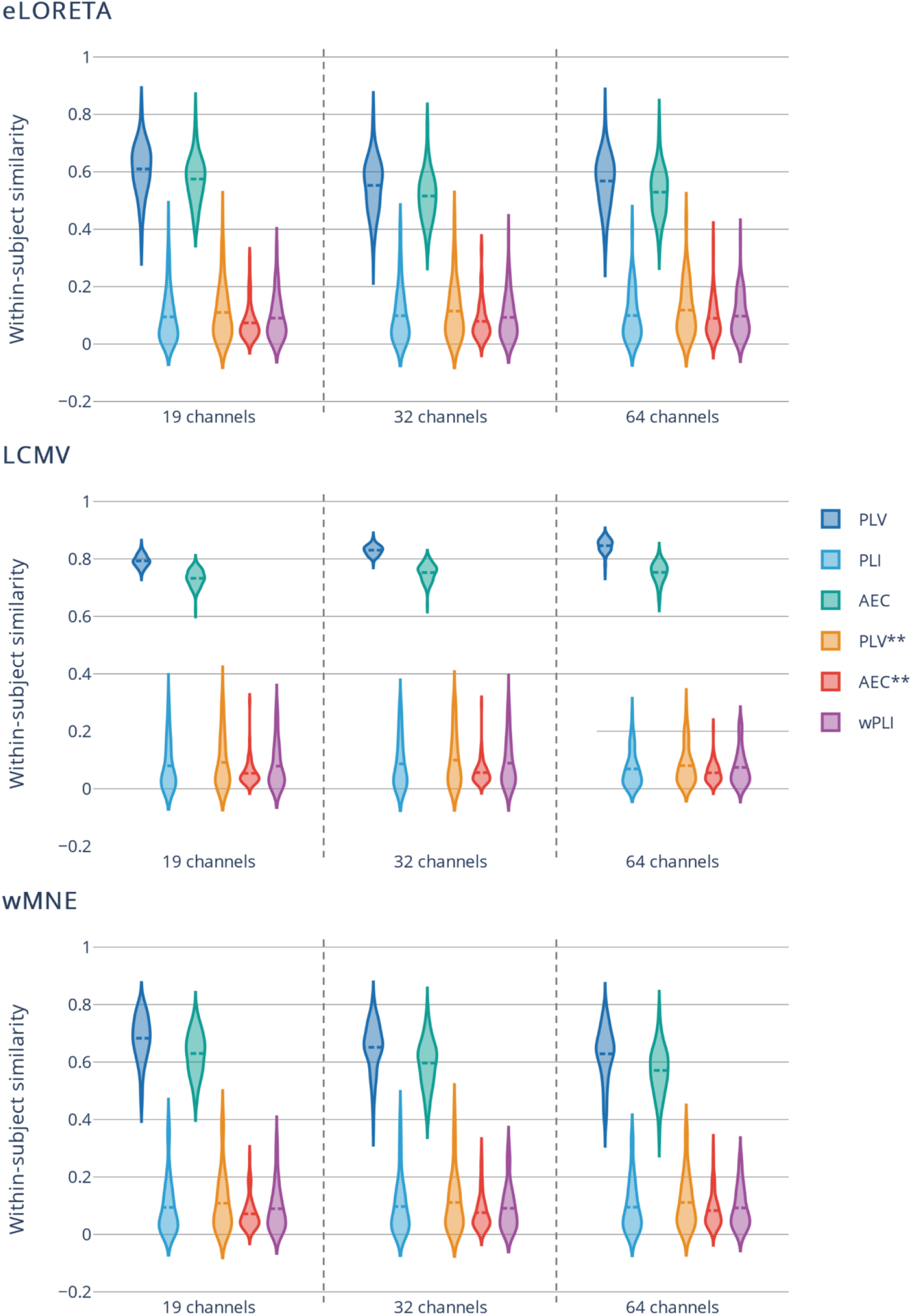
Within-subject similarity. For each condition (number of electrodes x inverse solution algorithm x connectivity measure), and for each subject, Pearson correlation was computed between connectivity matrices inferred from different epochs. All correlation values for a single subject were averaged to obtain a single value representing the degree of within-subject similarity/consistency. *eLORETA - exact low-resolution electromagnetic tomography. LCMV - linearly constrained minimum norm beamforming. wMNE - weighted minimum norm estimate. PLV - phase-locking value. PLI - phase-lag index. AEC - amplitude envelope correlation. wPLI - weighted phase-lag index. Asterisks (**) - source leakage correction*.

### 3.1. Effect of the number of electrodes

Group-level consistency, between-, and within-subjects similarity were investigated for different electrode densities (19, 32, 64 channels). Statistical tests showed a significant effect of the number of electrodes on the group-level consistency (F_(2, 5397)_=75, p<0.0001, R^2^=0.36) and between-subjects similarity (F_(2,206709)_=44, p<0.0001, R^2^=0.79), whereas no significant effect on the within-subjects similarity was detected (F_(2,4749)_=0.5523, p=0.6, R^2^=0.75). *Post-hoc* comparisons between group consistency values showed significant but very small differences between those obtained with 19 (0. 9484 ± 0. 0378) and 32 channels (0. 9524 ± 0. 0328) (p<0.01) and 19 and 64 (0. 9505 ± 0. 0351) channels (p<0.01). The difference between results obtained with 64 and 32 channels was not significant (p=1). Regarding the inter-subject similarity, all pairwise comparisons were significant (p<0.0001). Fig. 5 A) shows the distribution of consistency values at group-, inter-, and intra-subjects levels with respect to different channel densities (all source reconstruction algorithms and functional connectivity values are included). Fig. 2-5 do not show clear differences between different channel configurations in contrast to the results of statistical tests. However, these differences are more evident by taking into account the distribution of group consistency and between-subjects similarity values as shown in Fig. S4. in the Supplementary materials.

### 3.2. Effect of the inverse solution algorithm

In Fig. 5 B), we illustrated the distributions of the group level consistency, between-, and within-subjects similarity values with respect to different source reconstruction algorithms (all channels configurations and connectivity measures are included). Based on the statistical tests, the choice of the source reconstruction algorithm (eLORETA, LCMV, wMNE) had a significant effect on group consistency (F_(2,5397)_=1484, p<0.0001, R^2^=0.36), between-subjects similarity (F_(2,206709)_=14141, p<0.0001, R^2^=0.79), and within-subjects similarity (F_(2,4749)_=166, p<0.0001, R^2^=0.75). All pairwise *post-hoc* comparisons were statistically significant (p<0.0001). For the group consistency, the mean and standard values obtained with eLORETA (0. 9586 ± 0. 0291) were significantly higher (although the differences were small) than those obtained with LCMV (0. 9519 ± 0. 0370) and wMNE (0. 9409 ± 0. 0370), respectively. For between-subjects similarity, the values obtained for LCMV (0. 4520 ± 0. 3643) were higher than those obtained with eLORETA (0. 4209 ± 0. 2928) and wMNE (0. 3293 ± 0. 2421). Similarly, the within-subjects similarities values were 0. 2505 ± 0. 2342, 0. 3123 ± 0. 3402, 0. 2711 ± 0. 2655 for eLORETA, LCMV, and wMNE, respectively.

**Fig. 5.**
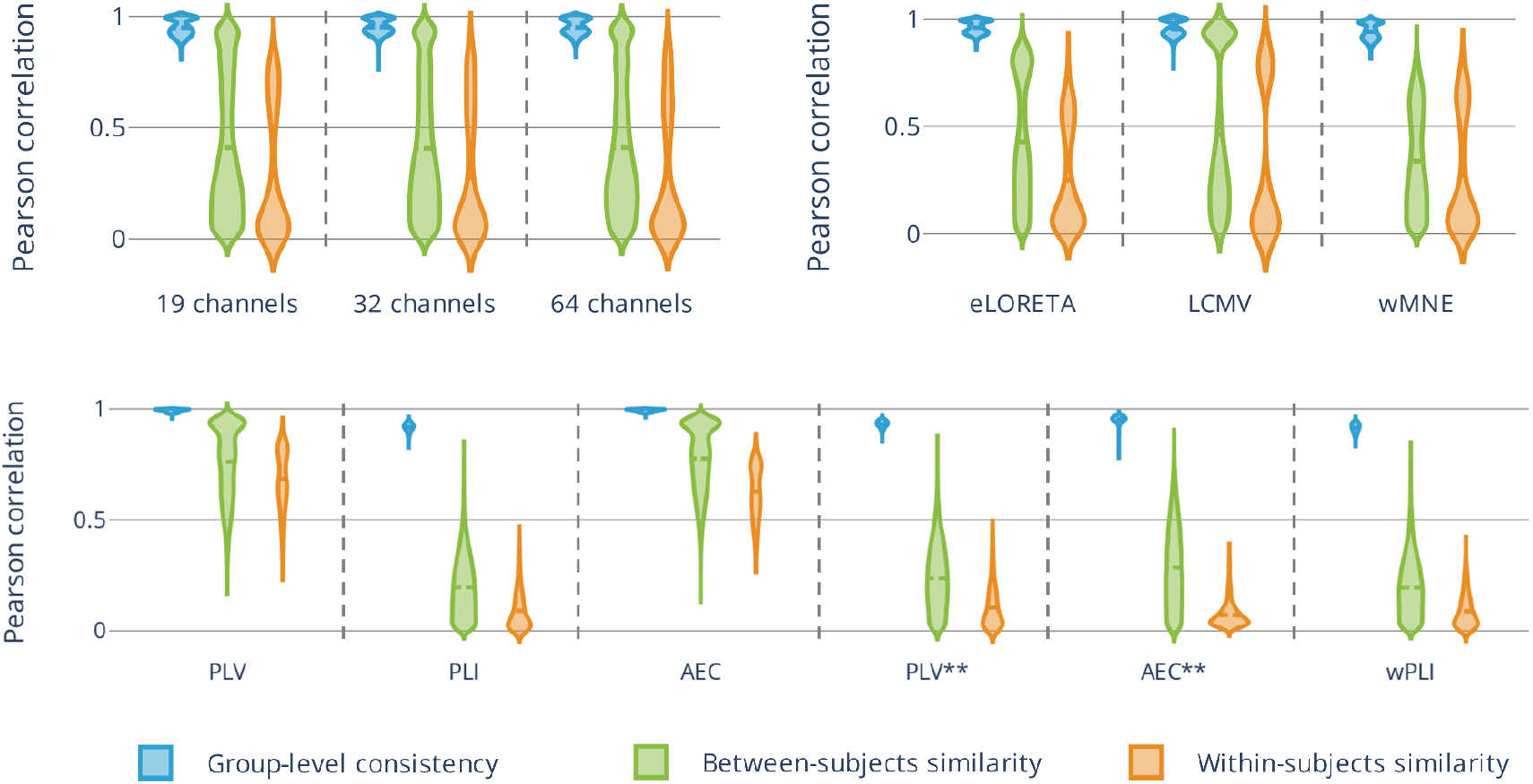
Violin plots of the group-level consistency, between-, and within-subjects variability with respect to A) different electrode configurations, B) source reconstruction algorithms, and C) functional connectivity measures. *eLORETA - exact low-resolution electromagnetic tomography. LCMV - linearly constrained minimum norm beamforming. wMNE - weighted minimum norm estimate. PLV - phase-locking value. PLI - phase-lag index. AEC - amplitude envelope correlation. wPLI - weighted phase-lag index. Asterisks (**) - source leakage correction*.

### 3.3. Effect of the connectivity measure

Group- and subject-level consistency values were both affected by the choice of the functional connectivity measure. Statistical tests showed a significant effect of the connectivity measure on group-level consistency (F_(5,5394)_=13874, p<0.0001, R^2^=0.36), between-subjects similarity (F_(5,206706)_=143027, p<0.0001, R^2^=0.79), and within-subject similarity (F_(2,4746)_=4951, p<0.0001, R^2^=0.75). At the group level, for all channel densities and source reconstruction algorithms, the highest consistency was obtained using PLV and AEC. No significant difference was observed between these two measures (p=0.97). Significantly lower consistency values were obtained in all methods that neglect zero lag (PLI and wPLI) or correct for source leakage (corrected AEC and PLV). Consistency values obtained with corrected AEC (0. 9539 ± 0. 0151), and corrected PLV (0. 9321 ± 0. 0154) were significantly lower than those with PLV (0. 9918 ± 0. 0067) and AEC (0. 9926 ± 0. 0057) (p<0.0001) and higher than those with PLI (0. 9178 ± 0. 0067) and wPLI (0. 9145 ± 0. 0208) (p<0.0001) who both had similar performance (p=1). A similar trend (higher values obtained with PLV and AEC as compared to other connectivity measures) was obtained when assessing between- and within-subjects similarity. Regarding inter-subject similarity, the mean value was 0. 7624 ± 0. 1631 and 0. 7768 ± 0. 1474 for PLV and AEC, 0. 2591 ± 0. 2159 and 0. 2305 ± 0. 1536 for corrected AEC and PLV, and 0. 1888 ± 0. 1428 And 0. 1868 ± 0. 1399 for PLI and wPLI, respectively. *Post-hoc* comparisons between different connectivity measures were all significant (p<0.0001; for PLV *vs* AEC p=0.001) except for PLI *vs* wPLI (p=0.99). Regarding intra-subject similarity, the mean value was 0. 6848 ± 0. 1283 and 0. 6284 ± 0. 1125 for PLV and AEC, 0. 1051 ± 0. 0925 0. 0905 ± 0. 0869 and and 0. 0709 ± 0. 0554 0. 0883 ± 0. 0742 for corrected PLV and corrected AEC, for PLI and wPLI, respectively. Significant statistical differences were obtained for all pairwise comparisons (p<0.0001; for corrected PLV *vs* wPLI p=0.016) except for corrected AEC *vs* PLI (p=0.9), corrected AEC *vs* wPLI (p=0.8), and PLI *vs* wPLI (p=0.2). In Fig. 5 C), we illustrated the distributions of the group level repeatability, between-, and within-subjects similarity values with respect to different functional connectivity measures (all channels configurations and source reconstruction algorithms). Results obtained for AEC and PLV with source leakage correction using the symmetric multivariate orthogonalization approach are shown in Fig. S5-S7 in the Supplementary Materials. It is noteworthy that the performance of those metrics were not stable across different channel densities.

## 4. Discussion

In neuroimaging research, the high dimensionality of data and the complexity of analysis workflows contribute to the issue of analytical variability, defined as the innumerable steps, parameters, and decisions often made arbitrarily within an analysis workflow. The issue is that the same data set could be analyzed, in many different ways, potentially leading to inconsistent results (Wicherts et al. 2016). Importantly, this makes research results harder to reproduce. In this work, we were interested in investigating the impact of analytical variability in the context of EEG-based brain functional connectivity analysis. We mainly focused on the effect of the choice of EEG electrode density, source reconstruction algorithm, and functional connectivity measure on the consistency/discrepancy in studies’ outcomes. In simulation-based studies, the availability of a ground truth allows an objective evaluation of different analyses, whereas, the lack of such ground truth poses a serious challenge when using empirical data. Therefore, alternative approaches are usually adopted with empirical data. For instance, we used group-level repeatability, inter-, and intra-subject similarities to assess the variability of the results induced by different analytical choices. Our objective was not to propose the ‘best’ analytical choices, but rather to initiate an effort of systematic quantitative investigation of the issue of analytical variability in estimating EEG-based cortical functional networks.

### 4.1. Effect of the number of electrodes

As established in (Srinivasan, Tucker, and Murias 1998), a high inter-electrode distance (i.e., corresponding to a low number of EEG electrodes) can induce aliasing, and therefore high spatial frequency signals are misrepresented as low spatial frequency signals due to the violation of the Nyquist criteria (*F_s_ > 2 × F_max_* (Srinivasan, Tucker, and Murias 1998; Song et al. 2015). In two previous simulation studies (Allouch et al. 2022, 2023), a significant improvement in the accuracy of network reconstruction was observed when increasing the number of electrodes (19, 32, 64, 128, 256). In this article, we tested the effect of three electrode configurations (64, 32, 19 channels) on the outcomes’ variability. In line with the reported studies, in this article, different channel densities resulted in significant (but small) differences in group-level repeatability, between-, and within-subjects variability.

### 4.2. Effect of the source reconstruction algorithm

Another critical influencing factor in the EEG source connectivity analysis is the algorithm chosen to reconstruct the cortical source activity. Several studies quantifying the performance of different inverse methods, in simulated and experimental EEG/MEG data, concluded that the choice of the inverse method significantly influences source estimation results (Anzolin et al. 2019; Mahjoory et al. 2017; Hedrich et al. 2017; Bradley et al. 2016; Grova et al. 2006; Halder et al. 2019; Tait et al. 2021). However, no consistent conclusions have been made regarding one method that would stand apart from the others in terms of performance. For example, (Anzolin et al. 2019) showed that LCMV had a better performance globally as compared to eLORETA. Similarly, in (Mahjoory et al. 2017), a relatively strong difference was found between LCMV beamformer on one hand, and eLORETA/wMNE solutions on the other hand. In (Bradley et al. 2016), the use of LORETA for source localization outperformed sLORETA and minimum norm least square. Results of source localization in (Halder et al. 2019) did not identify a clear winner between LCMV, eLORETA, MNE, and dynamic imaging of coherent sources (DISC). (Tait et al. 2021) summarized the conditions where each method can be recommended, following comparison of six inverse methods in resting-state MEG data. In line with the above-mentioned studies, we showed in (Allouch et al. 2022, 2023) that the choice of the source reconstruction algorithm has a significant effect on the accuracy of reconstructed cortical networks in the context of epileptiform and resting-state simulations. This same effect was also observed in this article: group-level consistency, between-, and within subjects similarity were all substantially affected by the choice of the source reconstruction algorithm.

### 4.3. Effect of the connectivity measure

The choice of the functional connectivity metric is also a critical step when reconstructing brain networks. A wide range of measures are used in the field, and each differs in the aspect of the data that is being investigated (see (Friston 2011; Pereda, Quiroga, and Bhattacharya 2005; Cao et al. 2022) for a review), resulting in significant variability of performance and interpretations (Colclough et al. 2016; H. E. Wang et al. 2014; Wendling et al. 2009; Hassan et al. 2017). For instance, (Colclough et al. 2016) assessed the consistency of different measures in experimental MEG resting state data and recommended using the correlation between orthogonalized, band-limited, power envelopes (AEC). On the other hand, following extensive simulation studies, (H. E. Wang et al. 2014) and (Wendling et al. 2009) both concluded that there is no ideal “one-fits-all” method for all data types: it was rather suggested to evaluate which conditions are appropriate for each method. In (Hassan et al. 2014) and (Hassan et al. 2017), in the context of epileptic spikes, wMNE combined with PLV had better accuracy as compared to other algorithms. In (Allouch et al. 2022, 2023), we tested several connectivity measures and obtained a significant variability in the accuracy of reconstructed cortical networks in the context of epileptiform and resting-state simulations (Allouch et al. 2022, 2023). In line with previous findings, significant differences in group-level consistency, between-, and within-subjects similarity were obtained for different connectivity measures tested in this work.

It is noteworthy that higher group-level consistency, between-, and within-subject similarity were obtained with PLV and AEC. In contrast, connectivity metrics that are supposed to be robust to source leakage and signal spread problems (PLI, wPLI, corrected PLV, and corrected AEC) resulted in substantially lower group-level consistency, between-, and within-subject similarity. Therefore, we tested whether high consistency values are correlated with source leakage and signal spread problems or not. As illustrated in Fig. S8 in Supplementary materials, there was no correlation between the contribution of each edge to the consistency value at the group level and edge length (i.e., the distance separating correspondent ROIs). The results shown were obtained using eLORETA as a source reconstruction algorithm and a montage with 64 electrodes. Similar results were obtained for other inverse solution algorithms and electrode configurations. Since there was no correlation between the contribution of each edge to the consistency values at the group level and the distance separating different ROIs, high group-level repeatability cannot be explained by source leakage and signal spread problem. The high group-level consistency values we obtained are therefore not a result of repetitive connectivity patterns induced by source leakage. In contrast, in (Colclough et al. 2016), the authors found that high group-level repeatability is spurious and caused by the signal spread problem. Moreover, the low within-subject similarity values we obtained are in line with the results obtained by (Fraschini et al. 2019), where methods correcting for the source leakage problem performed the worst in terms of within-subject network-based fingerprints. The authors proposed that metrics correcting for the source leakage problem (PLI, wPLI, corrected PLV, corrected AEC) may be obscuring individual network characteristics which results in lower within-subject epochs similarity.

### 4.4. Other possible sources of variability

The purpose of our paper was not to present an exhaustive investigation of all sources of analytical variability in the EEG source connectivity analysis. There exist numerous factors that were not addressed in this work and are thought to produce substantial variability in reported results. For instance, in (Šoškić et al. 2022), preprocessing and data cleaning steps have been shown to affect the outcomes in ERP analysis. Similar conclusions were obtained in (Clayson et al. 2021). In addition to the specific method chosen to process the data, the selected software package can be a substantial source of variability. This topic was more extensively tackled in fMRI as compared to EEG studies (Bowring, Maumet, and Nichols 2019; Glatard et al. 2015; Gronenschild et al. 2012). Recently, (Aya Kabbara et al. 2023) examined the degree of consistency between EEG software toolboxes used to preprocess evoked-related potentials. Other sources of variability in EEG source connectivity analysis are related to the inverse solution parameters (regularization parameter for example (Grech et al. 2008)), head models (Cho et al. 2015; Brodbeck et al. 2011; Wolters et al. 2006), and channels locations (Shirazi and Huang 2019). The definition of network nodes (voxel-wise networks, anatomical or functional atlases) and the corresponding spatial resolution are also critical factors resulting in significant discrepancies in results (Yao et al. 2015; de Reus and van den Heuvel 2013; Stanley et al. 2013; Fornito, Zalesky, and Bullmore 2010; Hayasaka and Laurienti 2010; Zalesky et al. 2010; J. Wang et al. 2009), in addition to the approach used to extract a single time series representative of an ROI (mean, principal component analysis…) (Zhou et al. 2009). Related to the functional connectivity computation, some sub-parameters are to be cautiously chosen such as epoch length (Fraschini et al. 2016; Wilson et al. 2015), and number of trials (Marquetand et al. 2019; Bastos and Schoffelen 2015). Finally, thresholding connectivity matrices (absolute *vs* proportional thresholds (van den Heuvel et al. 2017)) and whether to binarize them or not (Bassett and Bullmore 2017) are also debatable topics.

### 4.5. Possible solutions

The amount of analytical variability and its substantial impact on the results is a serious challenge for the neuroimaging community. To address this issue, several practices could be adopted. First, detailed documentation of the analysis is mandatory. However, with the emergent complexity of neuroimaging workflows, a detailed description of all analysis steps becomes exhausting, and even sometimes unachievable. In this case, the code itself becomes the most accurate documentation of the performed analysis (Niso, Botvinik-Nezer, et al. 2022). As a consequence, sharing underlying data, codes, and methods should become the norm to promote transparent analysis (Niso, Botvinik-Nezer, et al. 2022; Niso, Krol, et al. 2022; Pernet et al. 2020; Botvinik-Nezer et al. 2020; Poldrack et al. 2017; Wicherts et al. 2016; Bishop et al. 2015; Pernet and Poline 2015). Making the data and analysis available to the community fosters research replicability, and enables 1) running alternative analyses on the same data, and 2) validating the codes that were used (Botvinik-Nezer et al. 2020). To address the issue of p-hacking, pre-registration could be adopted to prevent the misuse of analytical variability and exploitation of the researcher’s degrees of freedom by setting analysis details prior to the study and prohibiting data-dependent choices. Although data and code sharing and pre-registration would not alleviate all the aspects of the analytical variability problem, it would make it transparent and quantifiable (Botvinik-Nezer et al. 2020). Another way to address the issue of analytical variability is to run many alternative analysis pipelines (and report them eventually) on the same data set (Botvinik-Nezer et al. 2020). This is becoming steadily more feasible with the increased automation of neuroimaging workflows. Such validation procedures could be, ideally, done by several teams. In this line, “multiverse analyses” have been proposed in other disciplines to increase analytic transparency (Steegen et al. 2016; Hall et al. 2022; Patel, Burford, and Ioannidis 2015; Simonsohn, Simmons, and Nelson 2019). This approach consists of running the whole (or a large) set of possible analytical combinations of choices and reporting the corresponding results. The advantages of a multiverse analysis are a presentation of the robustness or fragility of results across different analytical choices and an identification of critical choices highly affecting the outcomes. Finally, raising awareness about the topic of analytical variability and its substantial effects on reproducibility and sound neuroimaging research and practices is of the utmost importance. This can be promoted and disseminated through workshops, training (Bishop et al. 2015), and ongoing open discussions about possible solutions.

## Code availability

The codes that support the results of this study are available at https://github.com/sahar-allouch/var-EEG-FC. We used Matlab (Matlab 2018), Brainstorm toolbox (Tadel et al. 2011), Fieldtrip toolbox ((Oostenveld et al. 2011); http://fieldtriptoolbox.org), OpenMEEG (Gramfort et al. 2010) implemented in fieldtrip, Automagic toolbox (Pedroni, Bahreini, and Langer 2019) for EEG preprocessing, R (R Core Team 2020) for statistical analysis, and Seaborn (Waskom 2021) and Plotly (Plotly Technologies Inc 2015) for visualization.

## Data availability

The data that support the findings of this study are available upon request from the author [V. P.].

## Acknowledgments

The authors would like to acknowledge the Lebanese National Council for Scientific Research (CNRS-L), the Agence Universitaire de la Francophonie (AUF), and the Lebanese University for granting Ms. Allouch a doctoral scholarship. This work was also supported by the Labex Cominlabs project PKSTIM and by the Institute of Clinical Neuroscience of Rennes (Projects EEGCog and EEGNET3). The authors would also like to thank Campus France, Programme Hubert Curien CEDRE (PROJET N° 42257YA), and the Lebanese Association for Scientific Research (LASER) for their support.

