## Supplementary Materials for "Effect of analytical variability in estimating EEG-based functional connectivity"

### Materials and Methods

#### EEG pre-processing

EEG data were pre-processed using Automagic, a MATLAB-based toolbox. Pre-processing configuration is presented hereafter (all parameters definition can be found at https://github.com/methlabUZH/automagic/wiki/Configurations):

**Residual Bad Channel Detection**

- **High Variance Criterion (HVC) =** 20
- **Cut-off** = 80
- **Rejection Ratio =** 1/(512 Hz*10sec)
- **Minimum Variance Criterion (MVC)** = 1
- **High Pass Filter:** cut-off frequency = 45 Hz.
- **Low Pass Filter:** cut-off frequency = 0.1 Hz.
- The filter order will be estimated according to eeg_filtnew() of EEGLAB by default.
- **EOG regression** is checked (EOG channels = [65 66]).
- **ICLAbel** is checked (with temporary high pass filter (cut-off = 2 Hz).
- **Interpolation:** spherical
- Exclude ICLabel components
- **Probability thresholds for muscle artifacts:** [0.8 1]
- **Probability thresholds for eye artifacts:** [0.8 1]
- **Probability thresholds for heart artifacts:** [0.8 1]
- **Probability thresholds for line artifacts:** [0.8 1]
- **Probability thresholds for channel artifacts:** [0.8 1]
- **High Pass Filter:** cut-off = 0.1 Hz; (pop_eegfiltnew()).
- **Low Pass Filter:** cut-off = 45 Hz; (pop_eegfiltnew()).
- **Quality Rating:**
- **Overall Threshold (mV):** [20 25 30 35 40]
- **Time Threshold SD (mV):** [10 20 30 40 50]
- **Channel Threshold SD (mV):** [10 20 30 40 50]
- **Downsampling Rate:** 1

### Results

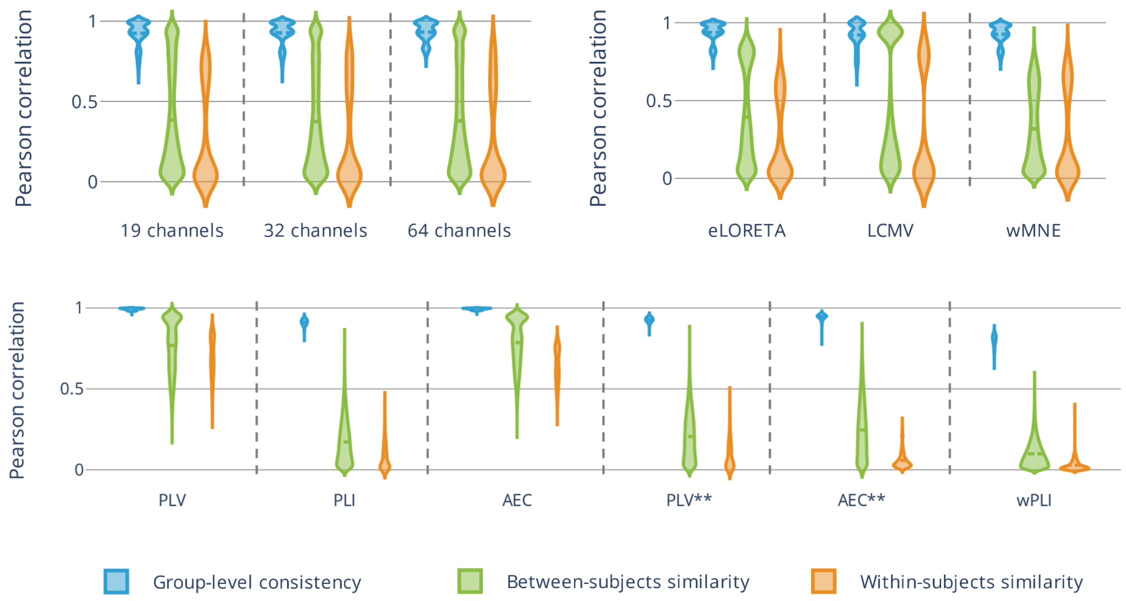


**Fig. S1.** Violin plots of the group-level consistency, between-, and within-subjects variability with respect to A) different electrode configurations, B) source reconstruction algorithms, and C) functional connectivity measures. Results correspond to the theta ([4-8]Hz) frequency band. *eLORETA - exact low-resolution electromagnetic tomography. LCMV - linearly constrained minimum norm beamforming. wMNE - weighted minimum norm estimate. PLV - phase-locking value. PLI - phase-lag index. AEC - amplitude envelope correlation. wPLI - weighted phase-lag index. Asterisks (**) - source leakage correction.*

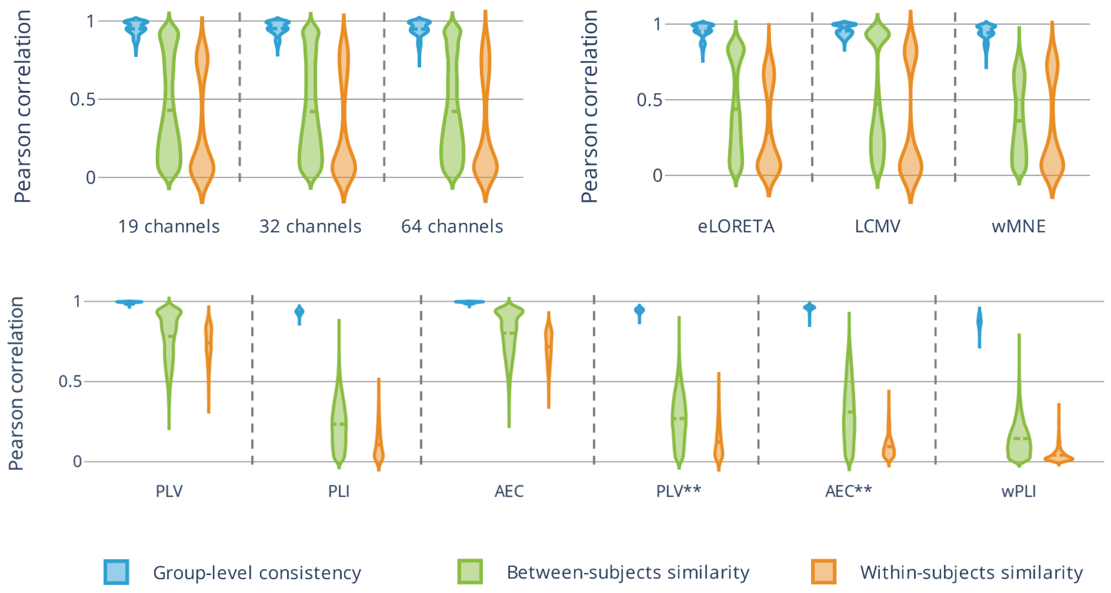


**Fig. S2.** Violin plots of the group-level consistency, between-, and within-subjects variability with respect to A) different electrode configurations, B) source reconstruction algorithms, and C) functional connectivity measures. Results correspond to the beta ([13-30]Hz) frequency band.

*eLORETA - exact low-resolution electromagnetic tomography. LCMV - linearly constrained minimum norm beamforming. wMNE - weighted minimum norm estimate. PLV - phase-locking value. PLI - phase-lag index. AEC - amplitude envelope correlation. wPLI - weighted phase-lag index. Asterisks (**) - source leakage correction.*

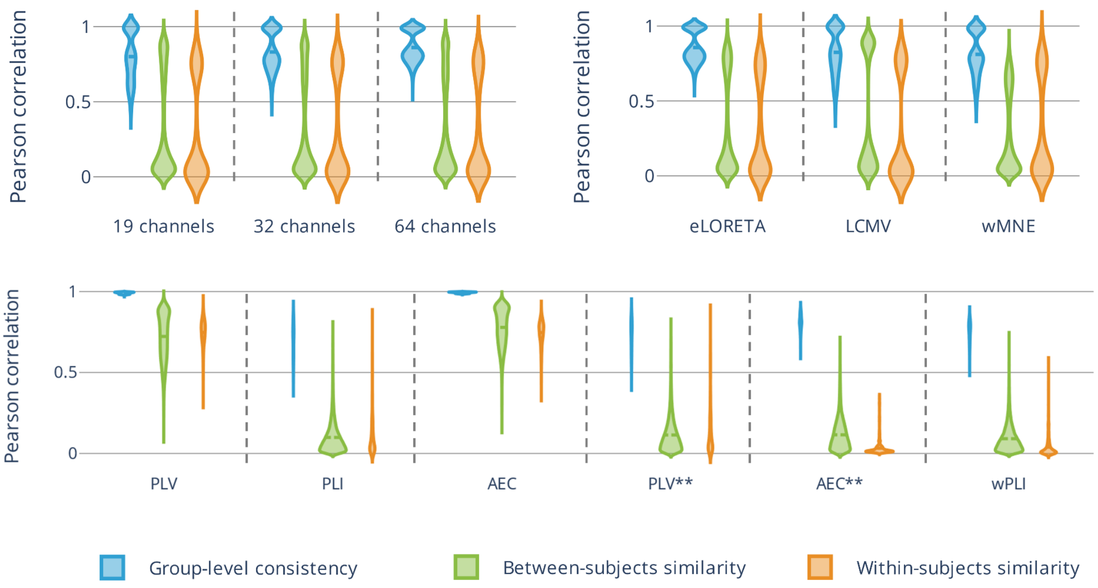


**Fig. S3.** Violin plots of the group-level consistency, between-, and within-subjects variability with respect to A) different electrode configurations, B) source reconstruction algorithms, and C) functional connectivity measures. Results correspond to the gamma ([30-45]Hz) frequency band. *eLORETA - exact low-resolution electromagnetic tomography. LCMV - linearly constrained minimum norm beamforming. wMNE - weighted minimum norm estimate. PLV - phase-locking value. PLI - phase-lag index. AEC - amplitude envelope correlation. wPLI - weighted phase-lag index. Asterisks (**) - source leakage correction.*

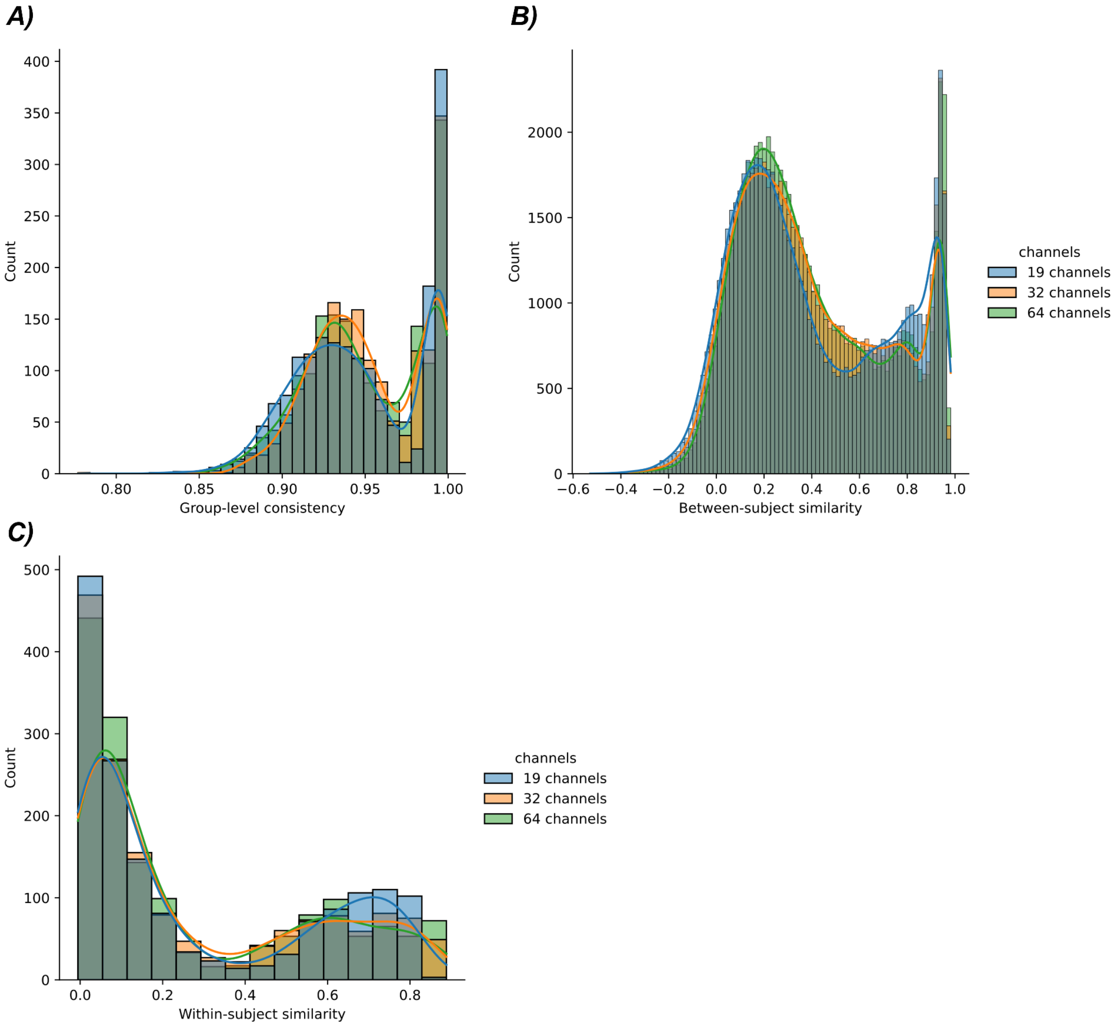


**Fig. S4.** Distributions of A) group-level consistency, B) between-, and C) within-subjects variability for different channel configurations (19, 32, and 64 electrodes). Significant differences obtained with statistical tests can be explained by the differences in the distribution plotted above (A-B).

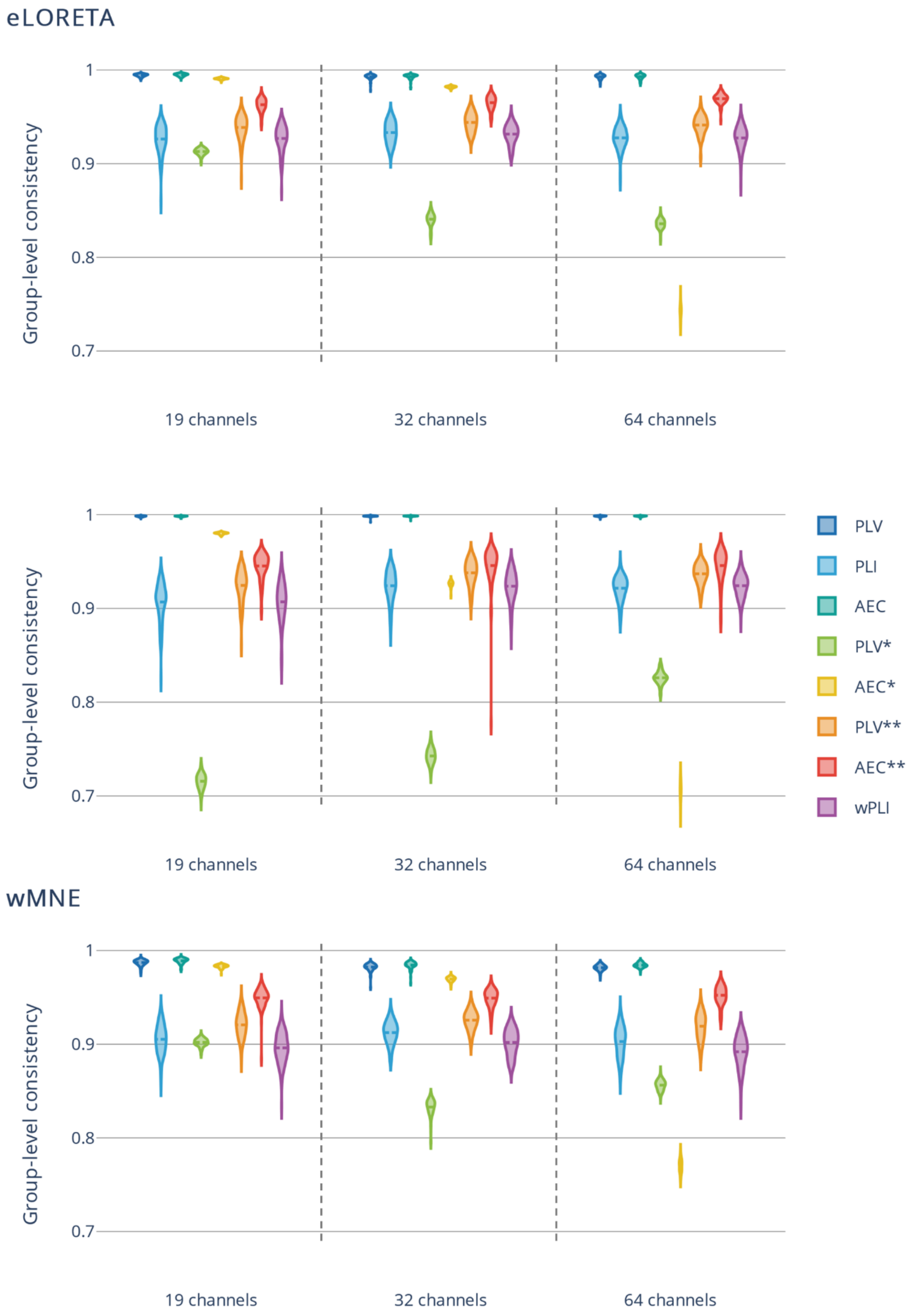


**Fig. S5.** Group-level consistency. The dataset was divided into two halves 100 times. For each condition (number of electrodes x inverse solution algorithm x connectivity measure), the Pearson correlation was computed between averaged connectivity matrices inferred from separate halves of the dataset. *eLORETA - exact low-resolution electromagnetic tomography. LCMV - linearly constrained minimum norm beamforming. wMNE - weighted minimum norm estimate. PLV - phase-locking value. PLI - phase-lag index. AEC - amplitude envelope correlation. wPLI - weighted phase-lag index. Asterisks: (*) - source leakage correction using the symmetric multivariate orthogonalization approach. (**) - source leakage correction using pairwise orthogonalization approach.*

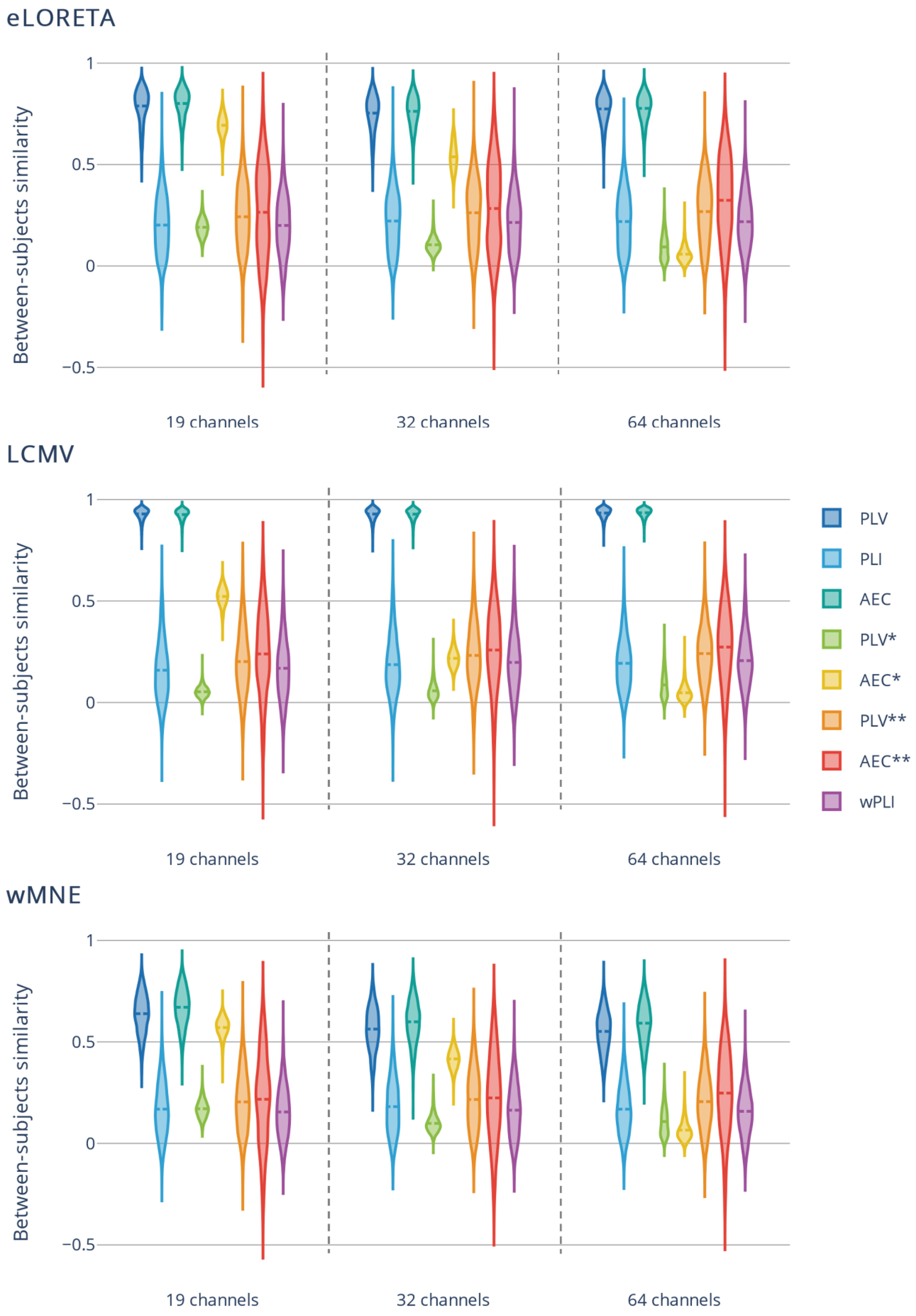


**Fig. S6.** Between-subjects similarity. Pearson correlation was computed between all subjects for each condition (number of electrodes x inverse solution algorithm x connectivity measure). *eLORETA* -

*exact low-resolution electromagnetic tomography. LCMV - linearly constrained minimum norm beamforming. wMNE - weighted minimum norm estimate. PLV - phase-locking value. PLI - phase-lag index. AEC - amplitude envelope correlation. wPLI - weighted phase-lag index. Asterisks: (*) - source leakage correction using the symmetric multivariate orthogonalization approach. (**) - source leakage correction using pairwise orthogonalization approach.*

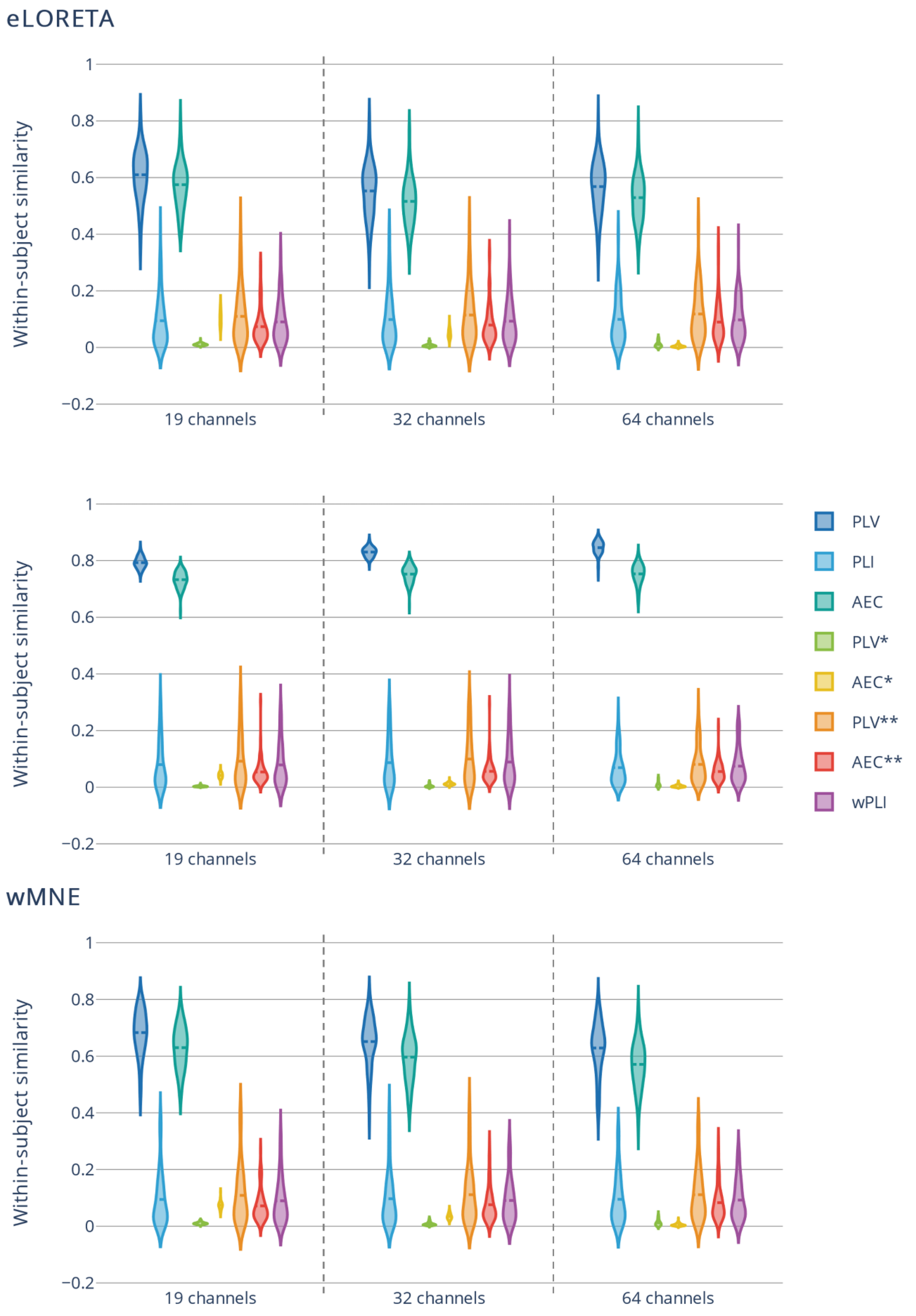


**Fig. S7.** Within-subject similarity. For each condition (number of electrodes x inverse solution algorithm x connectivity measure), and for each subject, Pearson correlation was computed between connectivity matrices inferred from different epochs. All correlation values for a single subject were averaged to obtain a single value representing the degree of within-subject similarity/consistency. *eLORETA - exact low-resolution electromagnetic tomography. LCMV - linearly constrained minimum norm beamforming. wMNE - weighted minimum norm estimate. PLV - phase-locking value. PLI - phase-lag index. AEC - amplitude envelope correlation. wPLI - weighted phase-lag index. Asterisks: (*) - source leakage correction using the symmetric multivariate orthogonalization approach. (**) - source leakage correction using pairwise orthogonalization approach.*

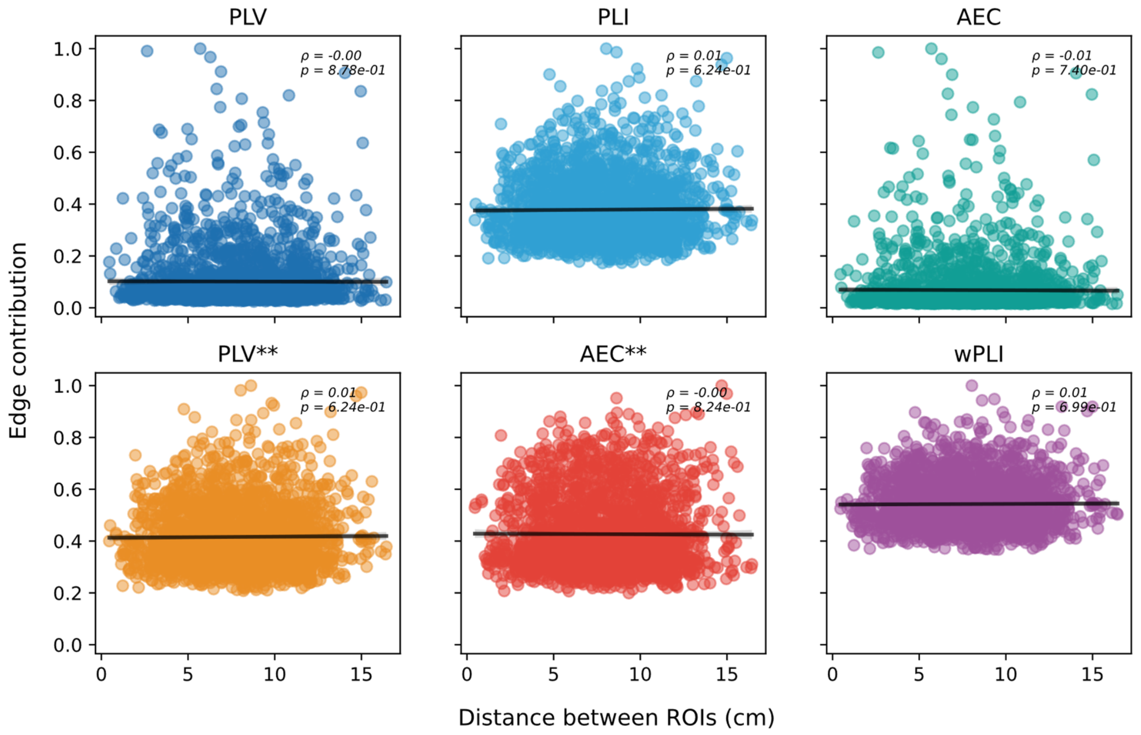


**Fig. S8.** Correlation between the edge contribution and the distance separating distinct ROIs for each connectivity measure (inverse solution algorithm = eLROETA, and number of electrodes = 64; similar result is obtained with different sensor density and source reconstruction algorithms). *PLV - phase-locking value. PLI - phase-lag index. AEC - amplitude envelope correlation. wPLI - weighted phase-lag index. Asterisk (**) - source leakage correction.*
